# Primed responses to damage signals mediate mycorrhiza-induced resistance in tomato plants

**DOI:** 10.1101/2024.08.01.606158

**Authors:** Zhivko Minchev, Juan M. Garcia, Estefania Pozo, Maria J. Pozo, Jordi Gamir

## Abstract

Arbuscular mycorrhizal fungi establish mutualistic associations with the roots of most vascular plants, enhancing plant immunity and activating mycorrhiza-induced resistance (MIR). In this study, we hypothesised that differential recognition of endogenous damage signals contributes to MIR in tomato plants. To test the hypothesis, we compared responses in mycorrhizal and non-mycorrhizal tomato plants after applying the cell-wall derived damage signal oligogalacturonides (OGs). We analysed the proteomic and metabolomic profiles, and the expression of marker genes related to plant defense, and the effects on plant resistance to the necrotrophic pathogen *Botrytis cinerea*. Our results show that mycorrhizal plants are more sensitive to these damage signals, as they respond to lower doses and exhibit stronger responses at the protein and metabolic level compared to non-mycorrhizal plants. Mycorrhizal plants showed primed accumulation of defense proteins, receptor kinases, flavonoids, and activation of the jasmonic acid and ethylene signaling pathways in response to OGs. Expression levels of the wall-associated kinase 1 (*slWAK1*) gene, coding for an OG receptor kinase in tomato, are elevated in mycorrhizal plants, and MIR against *B. cinerea* is abolished in a *wak1* mutant. Together, these results provide the first indication that self-damage recognition is essential to induce MIR against *B. cinerea*.

**Highlight:** Mycorrhizal tomato plants exhibit enhanced sensitivity to damage signals, leading to primed defense responses and induced resistance to fungal pathogens.

## Introduction

In nature, plants continuously interact with microorganisms, some potentially pathogenic and others neutral or even beneficial for plant health. The first ones can harm the plants using different strategies depending on their lifestyle. In contrast, beneficial microorganisms can establish mutualistic associations with the plants that benefit both partners (Edlinger et al., 2022). The plant immune system is central to control the interactions, facing the challenge of preventing infections by pathogens while promoting interactions with beneficials (Zhang and Kong, 2022). Root-associated beneficial microbes can improve plant nutrition, growth and stress tolerance, and they can boost plant immunity leading to induced resistance (IR) against pathogens and insects (Pieterse et al., 2014; De Kesel et al., 2021, Flors et al., 2024). Microbial-IR is associated to the priming of plant defenses through a fine-tuned regulation of plant immunity (Van der Ent et al., 2009, Flors et al., 2024). Different groups of root-associated microbes can trigger IR, being arbuscular mycorrhizal fungi (AMF) one of the most studied (Pozo and Azcon-Aguilar 2007, Jung et al., 2012; Pozo et al., 2021; Fiorilli et al., 2024). While some signalling processes regulating microbial-IR have been unveiled (Pieterse et al., 2014; Fiorilli et al., 2024), the early signals leading to primed defense activation are still unclear. Unravelling these early signals and events is essential for a better understanding and potential optimization of microbial-IR.

AMF are obligated biotrophs from the phylum Glomeromycota that establish symbiotic associations -known as arbuscular mycorrhizas (AM)-with the roots of about 80% of terrestrial plants. In the AM symbiosis the plant provides carbon-rich metabolites to the fungus, while the symbiosis improves plant acquisition of inorganic nutrients and water from the soil, and improves plant stress tolerance-resistance (Smith and Smith, 2011). In the onset of the interaction, during root colonization by the AMF, inducible responses are precisely locally activated and rapidly attenuated or modulated (Cameron et al., 2013, Martínez-Medina et al., 2019, Fiorilli et al., 2024). However, once the symbiosis is well-established, roots and leaves of mycorrhizal plants can display a primed defensive capacity (Sanmartín et al., 2020a; Rivero et al., 2021; Lidoy et al., 2024). Defense priming boosts basal defenses leading to IR with no major fitness costs (van Hulten et al., 2006; Martínez-Medina et al., 2016), and it is the main mechanism operating in mycorrhiza induced resistance (MIR). MIR refers to the enhanced resistance to pests and pathogens frequently observed in AMF-colonized plants, resulting from a primed defensive capacity associated to the mycorrhizal symbiosis (Pozo and Azcon-Aguilar, 2007; Jung et al., 2012; Cameron et al., 2013; Fiorilli et al., 2024). MIR against a broad spectrum of pathogens and herbivorous insects has been shown in many plant species, including diverse agronomically relevant crops such as melon (*Cucumis melo*), rice (*Oryza sativa*), peanuts (*Arachis hypogaea*), potato (*Solanum tuberosum*), tomato (*Solanum lycopersicum), and fruit trees as citrus and apples* (Campos-Soriano et al., 2012; Jung et al., 2023; Nair et al., 2015; He et al., 2017; Schoenherr et al., 2019; Sanmartín et al., 2020a; Rivero et al., 2021; Manresa-Grao et al., 2024; Minchev et al., 2024; Fiorilli et al., 2024).

Phytohormone signalling is central for defense priming, and in particular, priming of Jasmonic acid (JA)–dependent defenses play an essential role in MIR (Jung et al., 2012; Nair et al., 2015; Basso and Veneault-Fourrey, 2020; Lidoy et al., 2024). Along with JA-regulated defense responses, mycorrhiza primes alternative metabolic responses in systemic tissues. Flavonoids, terpenoids, lignans and other phenylpropanoid derivatives show primed accumulation in leaves of different plant species (Adolfsson et al., 2017; Hill et al., 2017; Sanmartín et al., 2020b). While these primed responses associated to the MIR display are being characterized, more research is needed to decipher the early signalling events triggering the boosted response associated to MIR.

Plant and animal immunity rely on monitoring systems that detect signals from the aggressors (“non-self” signals, including diverse microbe associated molecular patterns) and signals for self-damage recognition (Jones et al., 2024, Tanaka and Heil, 2021). Endogenous damage signals, commonly known as Damage-Associated Molecular Patterns (DAMPs), are endogenous molecules, with a specific function in intact cells, that are released to the apoplast upon cell disruption and perceived by adjacent intact cells as danger signals. DAMPs recognition triggers an alarm state in the plant that activates DAMPs-triggered immunity, activating early signalling responses regulating plant defenses (Galletti et al., 2008; Souza et al., 2017; Duran-Flores and Heil, 2017; Tripathi et al., 2018; Bacete et al., 2018; De Lorenzo and Cervone, 2022). DAMPs include sucrose, glutamate, nucleotides such as extracellular ATP and DNA as well as fragments of a disrupted cell wall like oligogalacturonides (OGs) and cellobiose (De Lorenzo et al., 2018; Jewell et al., 2019; Rassizadeh et al., 2021; Tanaka and Heil, 2021; Molina et al., 2024). The exogenous (non-host) and endogenous (host/self) danger signals (DAMPs) most likely act together to shape the appropriate immune response (Gust et al., 2017). However, there is still a big knowledge gap on how damage perception contributes to microbe-IR and specifically to MIR.

OGs are fragments from the cell wall pectin that are released to the apoplast due to the action of fungal or endogenous polygalacturonases after wounding (Gravino et al., 2015; Costantini et al., 2024). They act as DAMPs, triggering immune responses after their perception like Reactive Oxygen Species (ROS) production, MAP Kinase activation, callose deposition, upregulation of defense-related genes and production of phytoalexins and other defensive compounds, inducing pathogen resistance in different plant species (Ferrari et al., 2013; Pontiggia et al., 2020). The responses activated by DAMPs are tightly regulated by phytohormones; for instance, application of OGs to tomato plants induces JA, SA and ET signalling, and increase pathogen resistance (Gamir et al., 2020). The role of OGs in the primed defense responses in mycorrhizal plants is yet to be explored.

Wall-associated kinases (WAKs) are cell wall integrity sensors (Kohorn, 2016). In Arabidopsis, WAK1 is proposed as a receptor for OGs (Brutus et al., 2010) and is an essential contributor to disease resistance (Stephens et al., 2022). Wounding injury upregulates the expression of *WAK1* (Wagner and Kohorn, 2001, Gramegna et al., 2016) and *WAK1* overexpression increase plant resistance to the necrotrophic pathogens *Botrytis cinerea* and *Pectobacterium carotovorum* (Brutus et al.,2010; De Lorenzo et al., 2011). In tomato, WAK1 associates with flagellin receptors for full responsiveness to *Pseudomonas syringae* infection (Zhang et al., 2020) and the *Slwak1* mutant showed deficient responses to osmotic stress (Meco et al., 2020). However, the role of WAK1 in mediating OG signaling and the regulation of plant immunity remain unclear.

The present study aims to decipher whether DAMPs -more specifically OGs-recognition contributes to MIR. We used a multi-omics approach combined with targeted gene expression analysis to compare the responses to OG treatments in tomato plants colonized or not with the AMF *Funneliformis mosseae*. We addressed whether mycorrhizal colonization impacts plant responsiveness to OG perception at the metabolomic, proteomic and molecular level. Finally, we addressed the potential contribution of OG recognition to MIR by using the tomato mutant *Slwak1* (Meco et al., 2020). We tested the mutant’s responsiveness to OGs, its capacity to display OG-induced resistance and MIR against the necrotrophic foliar pathogen *B. cinerea*. Mycorrhizal plants responded to lower OG doses, showing enhanced sensitivity to OGs, and MIR was impaired in the *Slwak1* mutant. The results support that an enhanced sensitivity to perceive and respond to self-damage signals in mycorrhizal plants contribute to MIR.

## Material and Methods

### Plant and fungal material, growing conditions and experimental design

Seeds of wild type (WT) tomato (*Solanum lycopersicum*) cv Moneymaker and of a tomato Wall-Associated Kinase-1 (WAK1) mutant (described in Meco et al. (2020)) were kindly provided by the group of María C. Bolarín from the Department of Stress Biology and Plant Pathology, in the Centro de Edafología y Biología Aplicada del Segura (CEBAS-CSIC) from Spain.

The AMF *Funneliformis mosseae* BEG12 was maintained in an open pot culture of *Trifolium repens* mixed with *Sorghum vulgare* plants growing in a vermiculite-sepiolite (1:1) substrate in a greenhouse. The inoculum consisted of substrate containing infected root fragments, mycelia and spores (Rivero et al., 2018).

Tomato seeds were surface sterilized in 4% sodium hypochlorite, rinsed thoroughly with sterile water and incubated for 7 days in sterile vermiculite at 25°C. One week old plantlets were transferred to 300 ml pots containing a sterile mixture of sand:vermiculite (1:1). For mycorrhizal (Fm) treatments the growing substrate was mixed with 10% (v/v) of *F. mosseae* inoculum. For non-mycorrhizal (Nm) treatments, plants were irrigated with a filtrate (20 µm) of the *F. mosseae* inoculum to homogenise the general microbial populations, but free of AMF propagules. Plants were grown in a greenhouse with a 16 h light period, 70% relative humidity, and 26°C during the day and 18°C during the night. Once a week, tomato plants were watered with Long Ashton solution (Hewitt, 1996) with 25% of the standard phosphorus concentration. Plants grew for seven weeks before performing the chemical treatments and bioassays, to ensure a good establishment of the mycorrhrizal symbiosis.

### Determination of mycorrhizal colonization

Roots were sampled and washed at harvesting, cleared in 10% of KOH and AMF structures were stained in 5% ink in 2% acetic acid (García et al., 2020). To assess the percentage of root colonization by the AMF we used the grid-line intersection method (Giovannetti and Mosse, 1980) in a Nikon Eclipse 50i microscope under bright-field conditions.

### Oligogalacturonide treatments

Oligogalacturonides with a degree of polymerization between 10-15 were prepared as described in Benedetti et al. (2017). A solution of polygalacturonic acid (PGA) [2% (w/v; Alfa Aesar)] with endopolygalacturonase II [(0.1 RGU/mL) from *Aspergillus niger* Pectinase (Sigma)] was incubated in a water bath at 30°C for 180 min under shaking. The product is boiled for 10 min to inactivate the enzyme and put it at 4°C in an ice bath. The solution is mixed with cold 50 mM sodium acetate, 0.5 (w/v) PGA, and ice-cold ethanol, 17% (v/v). After O/N incubation the solution is centrifuged at 30,000 x g for 30 min. The pellet is solubilized and centrifuged again under the same conditions. The supernatant is dialyzed against ultrapure water [(cut-off of 1000 Da) (Spectra/Por®)] and lyophilized.

For elicitation, the fourth true leaf of seven weeks-old tomato plants were sprayed, using an aerograph, with an aqueous solution of OGs at two different concentrations, 5 µg/mL (Nm5/Fm5) or 50 µg/mL (Nm50/Fm50). Control plants (0 µg/mL; Nm0/ Fm0) were sprayed similarly with water. During the spray application of the selected leaf, the rest of the plant was covered with plastic to avoid contact with the solution. The systemic response to the treatment was analysed in the sixth fully developed –untreated-leaf, harvested 6 hours after the treatment, frozen in liquid nitrogen and kept at −80°C for further molecular analysis.

### DAB staining

Histochemical detection of H_2_O_2_ was performed using 3,3’-diaminobenzidine (DAB). Staining solution: 1 mg DAB/mL of distilled water in pH 3.3 using HCl 0.1 M. Tomato leaves sprayed with the OGs solution were immersed in DAB staining solution at the indicated time points after treatment. The tubes were conserved overnight in dark, and the next day the solution was replaced by Chloral hydrate (10 g / 4 mL distilled water) to remove the chlorophyll. The total number of pixels corresponding to plant leaves in micrographs was quantified using GIMP (2.6.12) software to estimate H₂O₂ content.

### Proteomics studies

Protein extraction was performed using Plant Total Protein Extraction Kit (sigma-Aldrich) following manufacturer’s instruction. Protein identification was carried out in the proteomic facility PROTEORED© of the University of Alicante (Spain) as detailed in Belchí-Navarro et al. (2019) with some modifications. In brief, 5 biological replicates per treatment (4 replicates in the case of Nm50 treatment) were extracted (30 mg) and digested with trypsin. Peptides were passed through a C18 column (Pierce® C18 Spin Columns, Thermo Scientific) and resuspended in formic acid (0.1%). Peptides were analysed with an Agilent 6550 iFunnel Q-TOF mass spectrometer (Agilent Technologies) coupled to an Agilent 1290 UHPLC chromatograph using an Agilent AdvanceBio Peptide mapping column (2.1 mm × 250 mm, 2.7 μm particle size, operated at 50°C). Analytes were eluted with a linear gradient of 3 - 40% ACN in 0.1% formic acid and with a constant flow of 0.4 mL/min. The LC-MS output files were loaded in Progenesis QI for Proteomics (Nonlinear Dynamics) v4.0 label-free analysis software and a protein quantification based on the MS peak intensity was performed. MetaboAnalyst 4.0 (Chong et al., 2018) was used to create the sparse partial least squares discriminant analysis (sPLS-DA) and heatmap analysis, and Venn Diagrams were plotted using Mapman (https://mapman.gabipd.org/).

### Sample extraction and UPLC-ESI-QTOF conditions

The full scan metabolomic analysis was performed as previously described in Gamir et al. (2014). Thirty milligrams of freeze-dried pulverized leaf were incubated for 30 min with 1 mL of extraction solution (MeOH: H2O (10:90) containing 0.01% of HCOOH) at 4°C. After the incubation, the product was centrifuged for 25 min at 14000 rpm at 4°C. The supernatant was filtered through 0.2 μm cellulose filters (Regenerated Cellulose Filter, 0.20 μm, 13 mm D. pk/100; Teknokroma) and conserved at −20°C until use. 20 µl of extraction solution was injected into an Acquity UPLC system (Waters, Milford, MA, USA) coupled to a hybrid quadrupole time-of-flight instrument (QTOF MS Premier). Simultaneous detection of ion signals and fragments of parental ions was performed by the mass detector. Three biological replicates were analysed per treatment. The samples were randomly injected. For chromatographic separation we used a UPLC Kinetex 2.6 μm particle size EVO C18 100 A, 50 x 2.1 mm (Phenomenex) and solvents and gradients were as described by Gamir et al. (2014).

### Full scan data analysis

Data files were transformed into .cdf using Databrigde software (Masslynx 4.1, Waters). Full scan data was processed using the XCMS algorithm (www.bioconductor.org; Smith et al., 2006) in software R (http://www.r-project.org/). Peaks areas were normalized to dry weights. The visual analysis, sPSL-DA and cluster analysis were performed with metaboAnalyst 4.0 (Chong et al., 2018). For Statistical analysis (nonparametric Kruskal–Wallis test; p < 0.05), adduct and isotope correction, filtering, exact mass mapping and metabolic pathway exploration we used Marvis suit 2.0 (Kaever et al., 2014). For metabolite identification, we used an internal database (more than 200 compounds) and online tools for exact mass and fragmentation spectra analysis, Metlin (http://www.masspec.scripps.edu) and Massbank (www.massbank.jp). Metabolic pathway annotation was based on exact mass identification and matching online fragmentation spectra.

### Gene expression analysis

Gene expression analysis was performed by qPCR. RNA from leaves was extracted using Tri-Sure (Bioline, London, UK) according to the manufacturer’s instructions. The RNA was treated with NZY DNase I (NZYtech, Portugal), purified through a silica column using the RNA Clean and concentratorTM (Zymo Research, USA) and stored at −80 °C until use. For the cDNA synthesis, we use 1 μg of purified total RNA using the PrimescriptTM RT master mix (Takara, Japan) according to the manufacturer’s instructions. We analysed six biological replicates per treatment and the gene expression was normalized against the housekeeping genes Actin (Solyc03g078400), elongation factor 1-alpha and beta-tubuline. The relative quantification of specific mRNA levels was performed using the comparative 2– Δ(ΔCt) method (Livak and Schmittgen, 2001). The gene-specific primers are shown in Table S1.

### Botrytis cinerea infection

*Botrytis cinerea* CECT2100 (Spanish collection of type cultures, Universidad de Valencia, Burjassot, Spain) grew in potato dextrose agar (Difco, Le Pont de Claix, France) plates, supplemented with freeze-dried tomato leaves. *B. cinerea* spores were collected from 3 weeks old plates and incubated in 0.5X potato dextrose broth (Difco, Le Pont de Claix, France) as previously described (Sanmartín et al., 2020a). The inoculation was performed by applying 6 µL drops (5 × 106 spores/ml) to detached tomato leaves. Two drops were applied per leaflet. The leaves were maintained in hermetically sealed boxes with 100% of humidity at 21°C in darkness. Necrotic lesions were evaluated after 5 days.

### Statistical analysis

We conducted a student’s t-test for the gene expression data using Microsoft office Excel. For the infection assay, primed proteins and DAB staining we conducted ANOVA and posthoc LSD analysis using Statgraphics Centurion XVI. All the metabolome profiling data were analysed using a Kruskal-Wallis analysis provided in MarVis suite 2.0.

## Results

### Mycorrhizal tomato plants respond to lower OG concentrations than non-mycorrhizal plants

To test whether mycorrhizal symbiosis enhanced plant sensitivity to damage signals, we performed a proteomic study comparting non-mycorrhizal (Nm) and mycorrhizal tomato plants colonized by the AMF *Funneliformis moseae* (Fm). Mycorrhizal symbiosis was confirmed, with 20 ±4 % of the root length colonized by the fungus. We sprayed the fourth true leaf in Nm and Fm plants with different OGs concentrations, 0 (control), 5 and 50 µg/mL. We analysed the systemic response six hours after treatment, comparing the proteomic profile of untreated upper leaves through untargeted proteomics. A sPLS-discriminant analysis overview of the proteome showed generally overlapping profiles among the different treatments (Figure 1A). However, a more detailed analysis comparing the effect of the different OG concentrations in mycorrhizal and non-mycorrhizal plants separately revealed that OGs application have a higher impact in mycorrhizal plants (Figure 1B). In the same line, heatmap analysis did not show effect of the lower OGs dose in non-mycorrhizal plants where this treatment (Nm5) grouped together with the control plants (Nm0) (Figure 1C). In contrast, mycorrhizal plants treated with the lower OG dose (Fm5) are grouped with the non-mycorrhizal plants treated with the higher dose (Nm50), revealing that at proteomic level, mycorrhizal plants are able to respond to lower amounts of OGs than non-mycorrhizal plants (Figure 1C).

**Figure 1.**
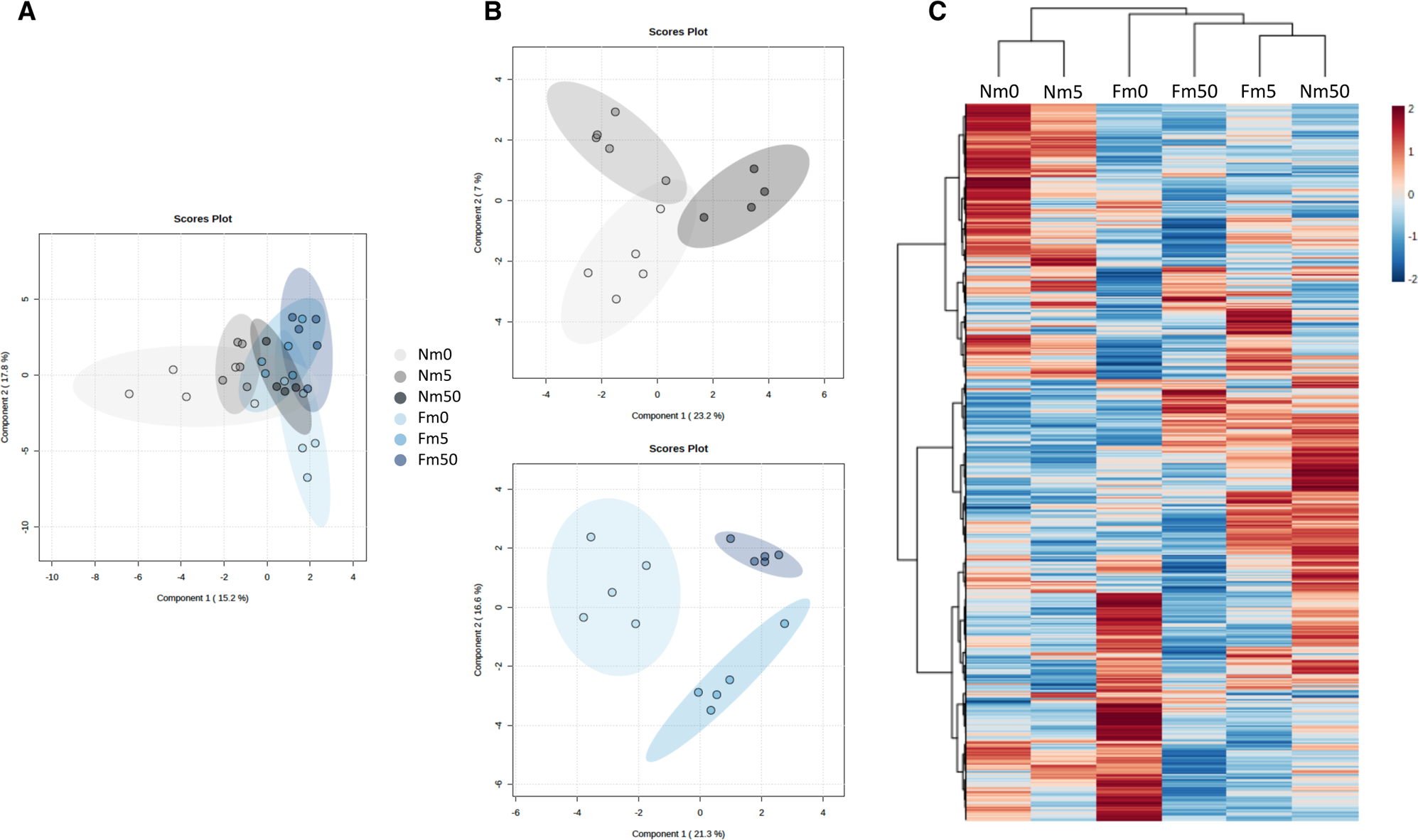
Proteomic analysis reveals that mycorrhizal plants are more sensitive to oligogalacturonides. (A) sPLS-discriminant analysis of the non-targeted proteomic study in leaves of non-mycorrhizal (nm) and mycorrhizal (fm) plants treated with different concentrations of OGs: 0, 5 and 50 µg/ml. (B) sPLSDA comparing the effect of OGs treatment in nm and fm plants separately. (C) Heatmap analysis of 200 proteins showing the highest changes among treatments according to ANOVA. Plants were colonized (Fm+) or non-colonized (Fm-) with the arbuscular mycorrhiza fungi Funneliformis mosseae and treated with two different oligogalacturonides (OGs) concentrations (5 and 50 µg/mL) or water as a control treatment (0 ug/mL). Each data point represents a biological replicate. All the samples were injected twice.

### Functional categorization of proteins with altered levels in mycorrhizal and/or OG-treated plants

To get a deeper insight into the effect of AMF and OGs in tomato, we performed a functional categorization of the differential proteins in the different conditions. First, we focussed on the differences associated to the mycorrhizal status of the plant by comparing the profile in water treated mycorrhizal and non-mycorrhizal plants. This study revealed 96 differential proteins, 30 of them being more accumulated and 66 proteins less accumulated in Fm0 than in Nm0 plants (Supplementary Figure 1A). Most of the differential proteins were related to the light reaction cycle, protein synthesis and degradation, RNA processing, abiotic/biotic stress and amino acid metabolism (Supplementary Figure 1B).

Regarding the response to OGs, the differential analysis showed 38 proteins differentially accumulated in non-mycorrhizal tomato plants treated with 50 µg/mL of OGs (Nm50) compared to water-treated plants (Nm0) (Supplementary Figure 2A). From them, 13 were more accumulated and 25 less accumulated after OG treatment. According to the functional categorization, the proteins with altered levels in Nm50 were related to light reaction, protein synthesis and degradation and amino acid metabolism, with most upregulated proteins not assigned to the major categories (Supplementary Figure 2B). Remarkably, the proteomic analyses revealed that AMF has a significant systemic impact on the leaf proteome, higher than the OGs treatment, while there is an overlap on the cell functions targeted by both treatments.

### Mycorrhizal symbiosis primes the accumulation of defensive related proteins in response to OGs

Defense priming is a major mechanism in MIR. We aimed to identify the primed responses to OGs in mycorhrizal plants. For that, we filtered the proteins accumulating to higher levels in mycorrhizal than in non-mycorrhizal plants in response to OGs. We identified 31 proteins with primed accumulation in response to 5 µg/mL of OGs and 20 primed proteins in response to 50 µg/mL of OGs (Figure 2). Strikingly, the primed response of mycorrhizal plants to 5 µg/mL or 50 µg/mL of OGs was completely different, with only one common primed protein identified in response to both concentrations, a root cap protein (solyc06g034130.2.1) (Figure 2). Among the mycorrhiza-primed proteins in response to OGs 5 µg/mL (Fm5), we identified a signalling receptor like kinase RLK (solyc04g074000.2.1), a cell wall pectin esterase (solyc06g009190.2.1), the abiotic stress related chaperon protein HtpG –also known as heat shock protein Hsp90-(solyc05g010670.2.1), and the biotic stress related proteins beta-1,3-glucanase (solyc01g008620.2.1), proteinase inhibitor 2 (Pin2, solyc11g021060.1.1) and endochitinase (solyc02g082920.2.1) (Figure 3A). Regarding the mycorrhiza-primed proteins in response to OGs 50 µg/mL (Fm50 treatment), we identified a serine threonine-protein kinase receptor (solyc10g005630.2.1), a glutathione peroxidase (solyc12g056230.1.1) and salicylic acid metabolism related pentatricopeptide repeat-containing protein (solyc07g025250.1.1) (Figure 3B). Therefore, an important part of the primed proteins in response to OGs are related to defense and signalling processes.

**Figure 2.**
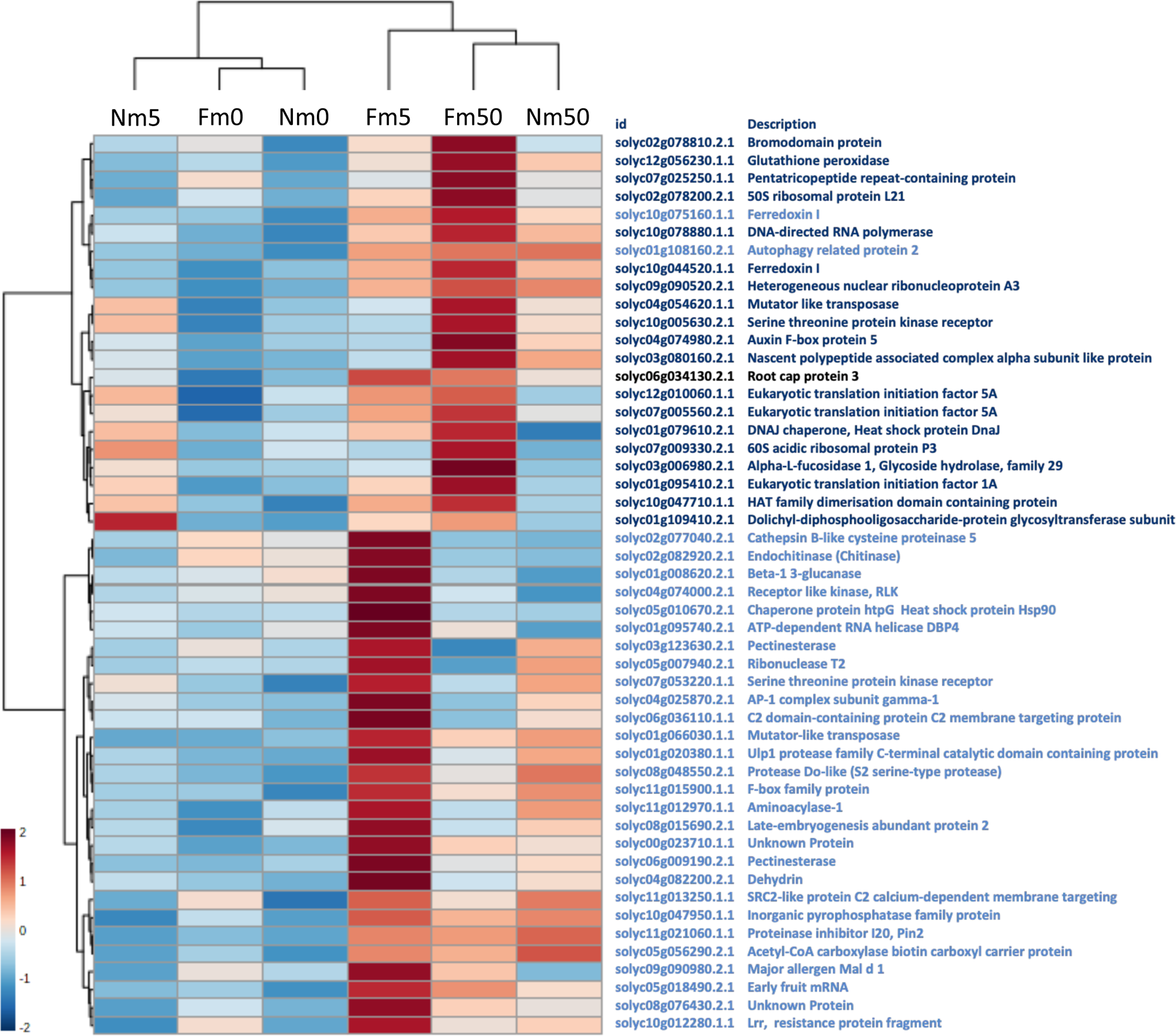
Heatmap representation of proteins. Heatmap representation displaying primed protein response in mycorrhizal plants (Fm) after treatment with OGs 5µg/ml (Fm5, green) or 50µg/ml (Fm50, blue) or both concentrations (black).

**Figure 3.**
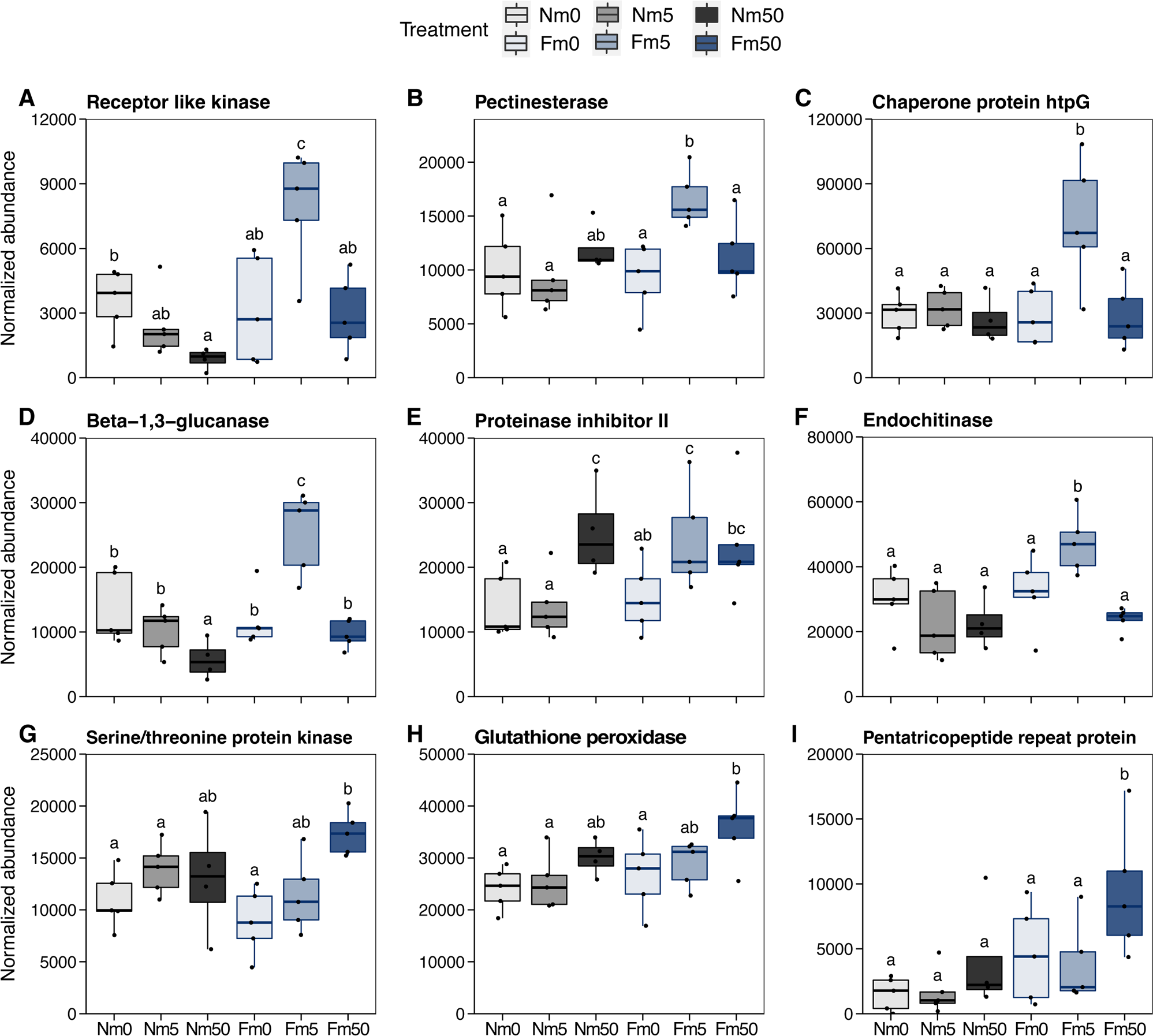
Proteins with primed accumulation in response to Oligogalacturonides. Boxplots of selected proteins with primed accumulation in response to (A) OGs 5µg/ml treatment and (B) OGs 50µg/ml. Treatments not sharing letters are significantly different based on ANOVA followed by LSD (p<0.05, n=5).

### Mycorrhizal plants display differential metabolic profiles upon OG application

To further explore the primed response to OGs in mycorrhizal plants, we performed an untargeted metabolomic analysis. The sPLS-DA showed that the application of 5 µg/mL of OG did not significantly impact the metabolomic profile in any of the plants. Regarding the higher OG dose, no major changes were observed in non-mycorrhizal plants after the treatment. In contrast, the application of 50 µg/mL of OGs leads to significant changes in the metabolic profile of mycorrhizal plants as compared to the water treated ones, confirming a primed response to these DAMPs also at the metabolic level (Figure 4a).

**Figure 4.**
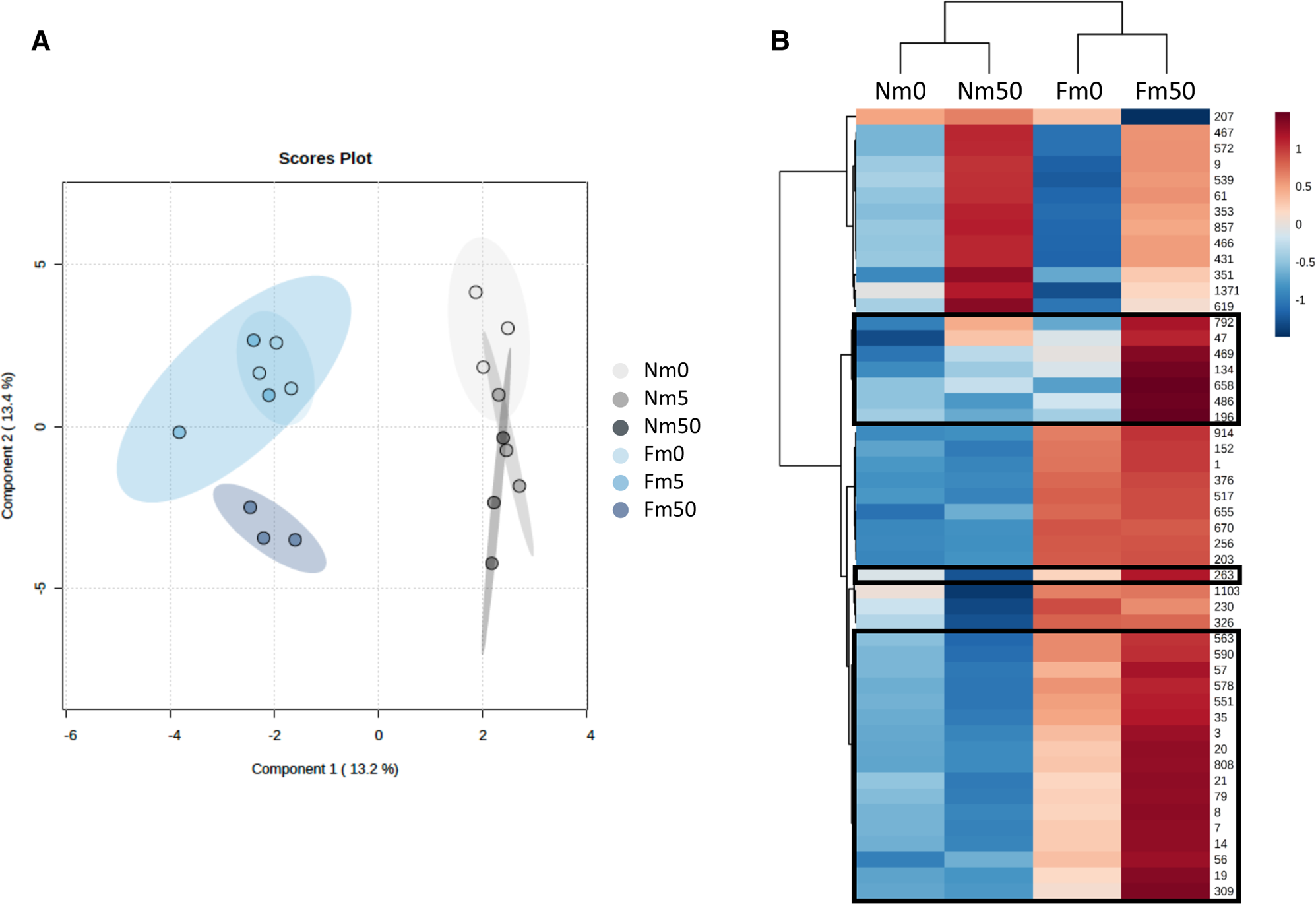
Mycorrhizal plants show primed metabolic responses to OGs. Non-targeted metabolomic profiling of systemic leaves responses in mycorrhizal (Fm) and non-mycorrhizal tomato plants (Nm-) treated with different OGs concentrations (0, 5 and 50 µg/mL). (a) The sPLS-DA of the signals indicates that only 50 µg/mL induces systemic responses in mycorrhizal plants. In non-mycorrhizal plants, OGs was unable to trigger differential metabolic responses. (b) Heatmap analysis of the signals showing significant changes among the selected treatments (p< 0,05, Kruskal-Wallis). Highlighted clusters contain primed signals.

Thus, we proceed with a more detailed comparative analysis of the samples treated with the higher dose (50 µg/mL). The heatmap analysis confirmed striking differences in the response to the OG treatment in mycorrhizal and non-mycorrhizal plants (Figure 4b). There is a set of compounds that are induced by the OG treatment in Nm plants that showed lower basal and induced levels in mycorrhizal plants. Besides that, three different clusters contained signals with a priming profile: higher accumulation in mycorrhizal plants in response to the treatment. Cluster 1 shows compounds strongly accumulated only in Fm50. Cluster 2 contains only one signal, repressed by OG in Nm and induced in Fm. Finally, cluster 3 corresponds to signals that show already elevated levels in water treated mycorrhizal leaves but are strongly induced upon OG treatment. From those signals we could identify four flavonoids, one amino sugar nucleotide and one anthocyanin, indicating that flavonoids could mediate primed responses in mycorrhizal plants after damage perception (Table S2).

### Mycorrhizal colonization primes transcriptional upregulation of JA- and Ethylene-signalling pathways in response to OG

To extend our findings, we analysed the expression levels of different genes related to plant immunity. We monitored the expression of genes related to JA and ET biosynthesis, signalling and response. OG treatment increased the expression of the JA-responsive genes Pin2 and LapA, at any of the concentrations tested (5 and 50 µg/mL) (Table 1) in both, mycorrhizal and non-mycorrhizal plants, although the induction of LapA was higher in Fm than in Nm plants upon the higher dose treatment. Interestingly, the transcript levels of the gene coding for multicystatin (*MC*) and polygalacturonase inhibitor protein (*PGIP*) showed a primed response in mycorrhizal plants treated with 5 and 50 µg/mL of OGs respectively. Specifically, *MC* gene expression in OG-treated mycorrhizal plants was 11 times upregulated as compared to water-treated non-mycorrhizal plants, whereas *PGIP* showed a 1.6-fold induction in OG-treated mycorrhizal plants. Finally, the expression levels of *GluB*, a defense-related, ET responsive gene, did not show any significant induction after OG treatment.

**Table 1.**
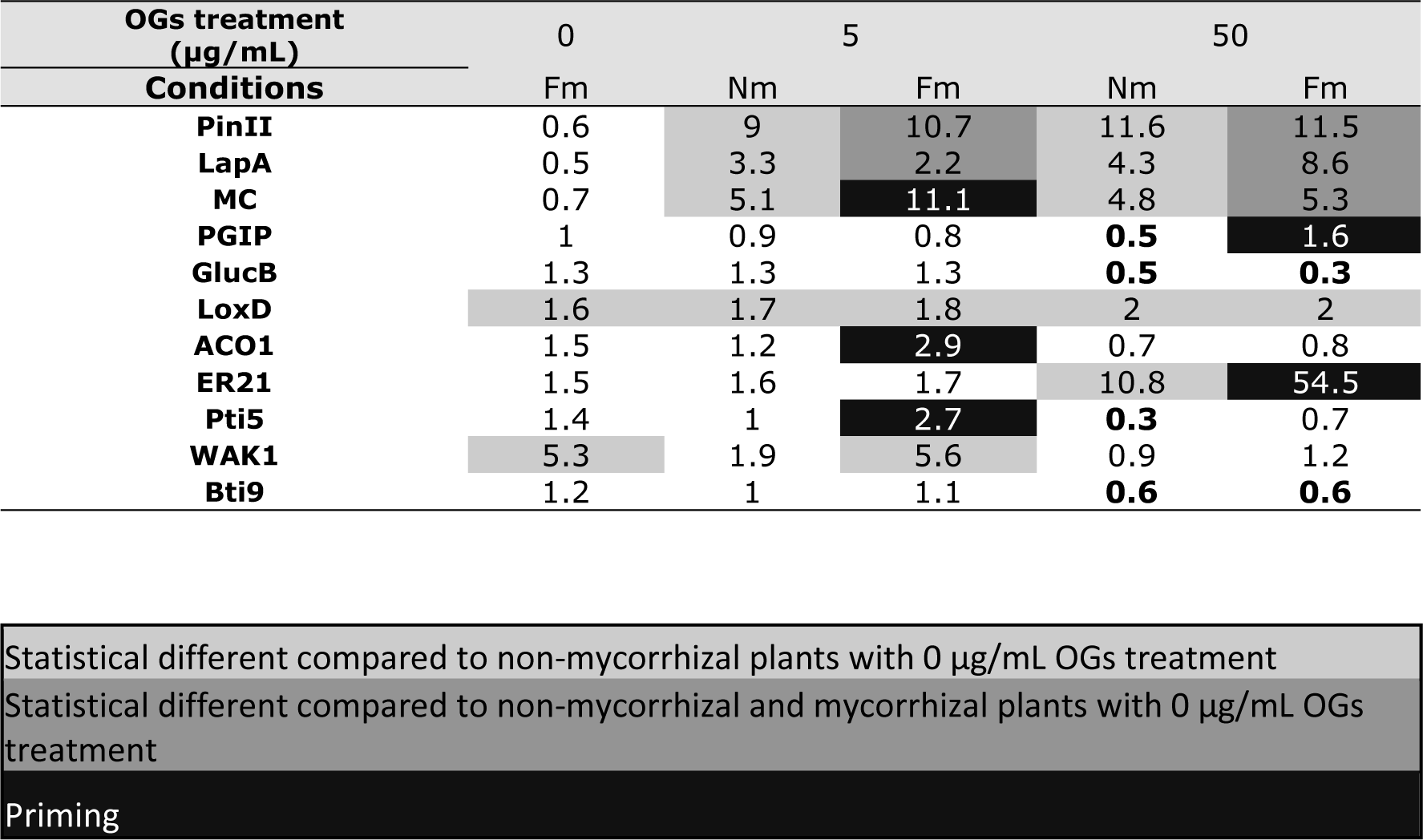
Gene expression analysis in mycorrhizal plants treated with OG. Quantitative RT-qPCR analysis of the expression levels of defence-related, biosynthesis, signalling and receptor genes. Non-colonised (Nm) tomato plants or colonised by the AM fungi Funneliformis mosseae (Fm) were treated with water, as a control treatment (0 µg/mL), or two different OGs concentrations (5 and 50 µg/mL). The values are normalized to control plants levels (Nm0). Relative quantification was performed using the comparative 2–Δ(ΔCt) method. Greyscale indicates significant differences to Nm0 (light grey), to both Nm0 and Fm0 (grey) and primed induction –response to OG treatment significantly different in Fm than in Nm plants-(dark grey). Bold numbers indicate repressed genes. (t-test; p-value<0.05; n=6).

| OGs treatment<br>(µg/mL) | 0 | 5 |  | 50 |  |
| --- | --- | --- | --- | --- | --- |
| Conditions | Fm | Nm | Fm | Nm | Fm |
| PinII | 0.6 | 9 | 10.7 | 11.6 | 11.5 |
| LapA | 0.5 | 3.3 | 2.2 | 4.3 | 8.6 |
| MC | 0.7 | 5.1 | 11.1 | 4.8 | 5.3 |
| PGIP | 1 | 0.9 | 0.8 | 0.5 | 1.6 |
| GlucB | 1.3 | 1.3 | 1.3 | 0.5 | 0.3 |
| LoxD | 1.6 | 1.7 | 1.8 | 2 | 2 |
| ACO1 | 1.5 | 1.2 | 2.9 | 0.7 | 0.8 |
| ER21 | 1.5 | 1.6 | 1.7 | 10.8 | 54.5 |
| Pti5 | 1.4 | 1 | 2.7 | 0.3 | 0.7 |
| WAK1 | 5.3 | 1.9 | 5.6 | 0.9 | 1.2 |
| Bti9 | 1.2 | 1 | 1.1 | 0.6 | 0.6 |

|  |
| --- |
| Statistical different compared to non-mycorrhizal plants with 0 µg/mL OGs treatment |
| Statistical different compared to non-mycorrhizal and mycorrhizal plants with 0 µg/mL OGs treatment |
| Priming |

Regarding JA and ET biosynthetic genes, the JA biosynthetic gene *LOXD* was induced by the OG treatment in both mycorrhizal and non-mycorrhizal plants. In contrast, the ET biosynthetic gene *ACO1* showed a primed response after the application of 5 µg/mL of OGs (Table 1). The transcript levels in OG-treated mycorrhizal plants were 2.9 higher than in control plants. However, the application of a higher dose did not change its expression compared to control plants, suggesting a dose dependent effect. Two ethylene (ET) responsive transcription factors, *ER21* and *PTI5* showed a strong primed response in mycorrhizal plants treated with 50 and 5 µg/mL of OG respectively. *ER21* levels showed a 54-fold change in mycorrhizal plants treated with 50 µg/mL compared to control levels. *PTI5* levels showed almost a 3-fold change in mycorrhizal plants treated with 5 µg/mL compared to control levels (Table 1). To summarize, these data show a primed activation in mycorrhizal plants of ET biosynthesis and signalling and primed upregulation of the JA responsive genes (*MC* and *PGIP*) in response to OGs.

Finally, we also analysed the expression levels of the OGs receptor, *WAK1* and the chitin receptor *Bti9* (Table 1). *Bti9* transcript levels remain unaltered by any of the treatments. Strikingly, the expression levels of *WAK1*, was 5-fold higher in water treated mycorrhizal plants compared to non-mycorrhizal plants. This induction was maintained after OG treatment with the lower dose, but not after treatment with the higher dose (Table 1). These higher basal levels in mycorrhizal plants may explain the enhanced responsiveness of those plants to the treatment.

### Oligogalacturonides perception is required for MIR against *Botrytis cinerea* in tomato

To test the relevance of the primed response of mycorrhizal plants to OGs in MIR we aimed to test MIR in a background deficient in OG perception. We first tested the OG response in the tomato mutant *wak1*, as it should show deficient activation of OG-triggered downstream signalling responses. Tomato *wak1* (Meco et al., 2020) contains a T-DNA insert in the promotor region of the gene that is homologous to the Arabidopsis WAK1, confirmed to be the OG receptor in this plant (Brutus et al., 2010). OGs are known to trigger oxidative burst in plants (Bellincampi et al., 2000; Galletti et al., 2008). We compared the OG response in the wild type and *wak1* mutant by analysing the OG-triggered hydrogen peroxide in both genotypes. DAB staining of the accumulated H_2_O_2_ revealed a weaker response in the mutant, as the wildtype Moneymaker accumulates more hydrogen peroxide than *wak1* after OG application (Figure 5a and b).

**Figure 5.**
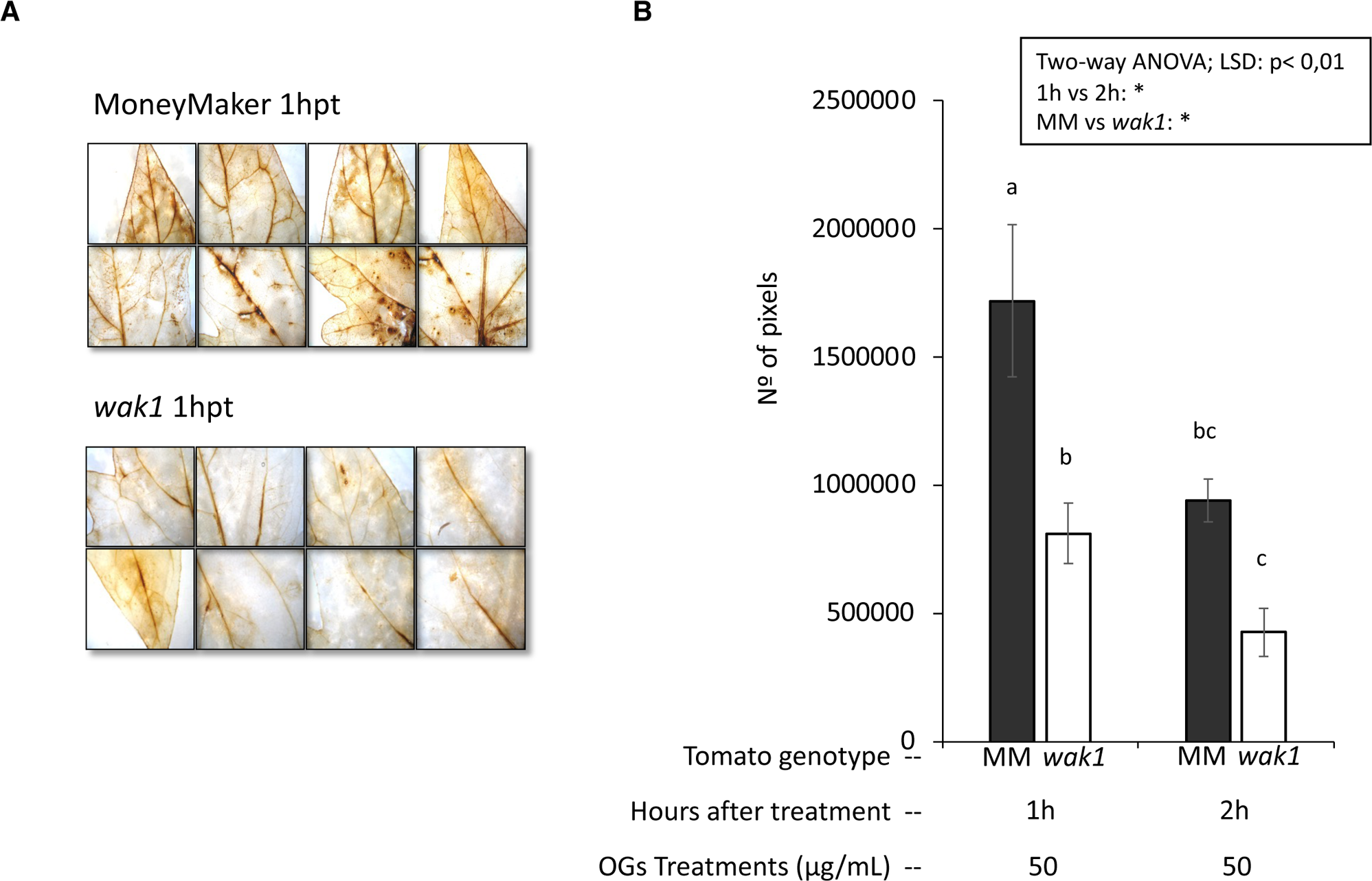
The OG receptor mutant, wak1, display a compromised response to OGs. Hydrogen peroxide accumulation in response to OG treatment in wak1 and wild-type tomato lines. Leaves of both lines were sprayed with a water solution of 50 ug/mL of OGs and harvested 1 and 2 hours after treatment (hpt). A) Illustrative images of DAB staining 1hpt. B) Quantification of the number of reddish-brown pixels corresponding to H2O2 accumulation. Bars show the means and the standard errors. Asterisks indicate significant factors (ANOVA: p<0,05; n=15-20); different letters indicate significant differences (LSD; p<0.01).

We hypothesised that the primed responses in mycorrhizal plants leading to MIR are dependent on OG signalling. To test it, we performed a bioassay to test the effect of mycorrhization on the resistance against the necrotrophic pathogen *B. cinerea* in the wildtype and *wak1* mutant lines. Mycorrhizal colonisation was well established in both lines (28 and 30% in wt and *wak1* mutant, respectively, with no significant differences among them according to t test, p>0.05). The plants were treated with different OG concentrations (0, 5 and 50 µg/mL). As in the previous experiment, we sprayed the plants with the OG solution in the fourth truly developed leaf and inoculated the pathogen in the sixth, untreated leaf six hours later. In the wild type, the mycorrhizal treatment and the OG50 treatment significantly reduced *Botrytis* infection (Figure 6). No synergistic effect was observed, since the mycorrhizal treatment alone reached the lower disease levels, similar to those in OG treated plants. In contrast, OG treatment failed to induce resistance in the *wak1* mutant, confirming the deficient responsiveness of the mutant to OG (Figure 6). In agreement with our hypothesis, no differences were observed in the resistance of mycorrhizal and non-mycorrhizal plants in the *wak1* mutant (Figure 6). Thus, OG-IR and MIR against *B. cinerea* were abolished in the mutant line, confirming our hypothesis. The results support our hypothesis, as proper OG perception is required to induce MIR against *B. cinerea* in tomato.

**Figure 6.**
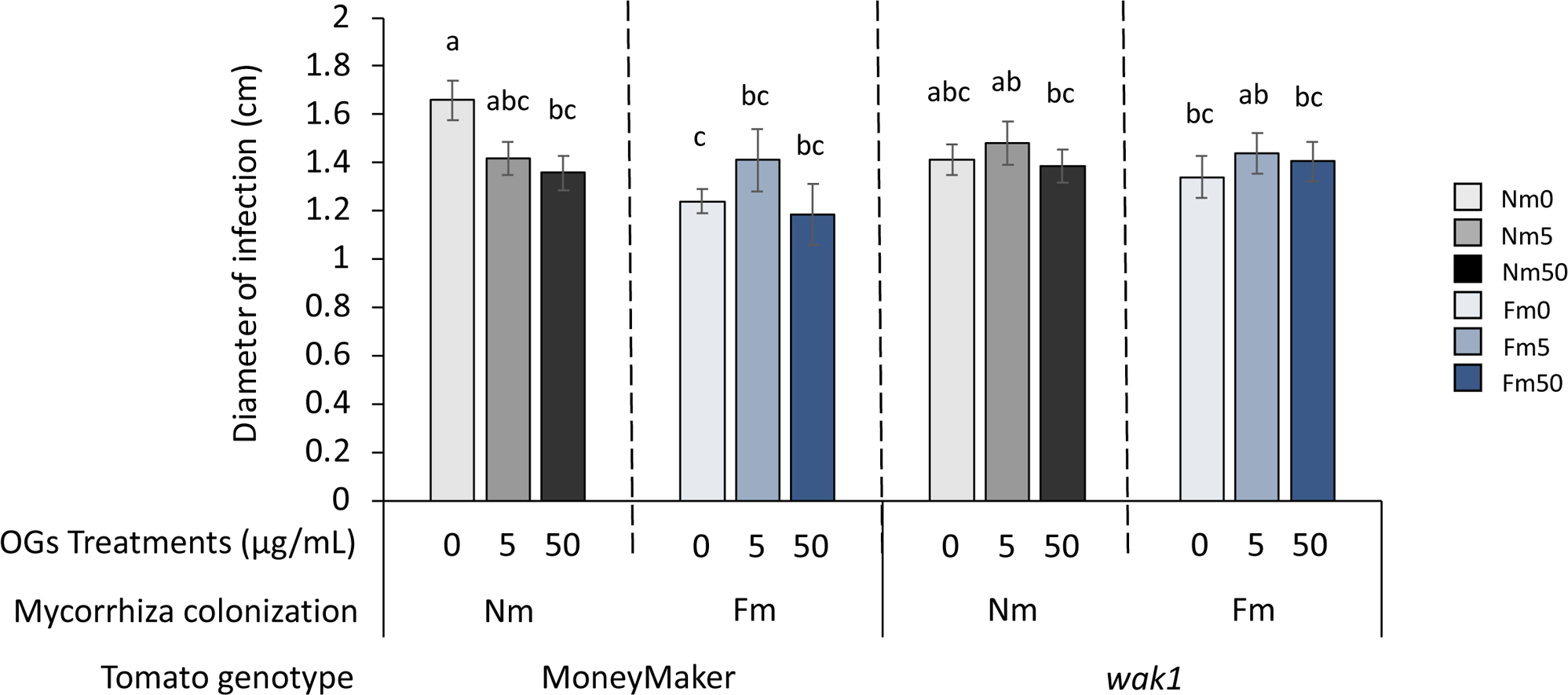
Tomato plants require a functional WAK1 to develop Mycorrhiza-IR (MIR) against *B. cinerea*. OG treatment and mycorrhizal inoculation significantly reduced B. cinerea lesion development in the wildtype genotype. The tomato mutant wak1 do not show either MIR or OG-IR against *B. cinerea*. Bars show the means and the standard errors of the diameter of the necrotic lesions in the leaves from non-mycorrhizal (Nm) and mycorrhizal plants with the AM fungus *Funneliformis mosseae* (Fm). The plants were treated with different oligogalacturonides (OGs) concentrations (5 and 50 µg/mL) or water as a control treatment (0 µg/mL) and drop inoculated with *B. cinerea* 6 hours post treatment. Different letters indicate significant differences (ANOVA: LSD; p<0.05; n=15-20)

## Discussion

Plant interaction with certain beneficial microbes can lead to a systemic protection against pests and diseases. This microbial-IR is generally associated to primed immune responses in the plant. While it is well recognized that self-damage recognition is an important part of the plant immune system, the contribution of damage signalling to IR remains unexplored. The overall goal of this study was to untangle the potential contribution of self-damage recognition to MIR. We show that treatments with OG as damage signals induce significant modifications in the proteomic and metabolomic profile of tomato plants, that differ among mycorrhizal and non-mycorrhizal plants. AMF colonization primes metabolic, proteomic and JA and ET related transcriptional responses to OGs, as mycorrhizal plants were able to respond to lower doses of OGs. Finally, we demonstrate that mycorrhizal tomato plants upregulated the expression of the *Slwak1* gene, likely coding for the OG receptor, and a *Slwak1 knock down* mutant was impaired in MIR against the necrotrophic pathogen *B. cinerea*. Together, these results support that DAMPs perception plays a relevant role in MIR, and highlight the relevance of self-damage recognition in the priming of plant immune responses by mycorrhiza.

The proteomic and transcriptomic profiles of mycorrhizal and non-mycorrhizal plants in response to OGs application revealed that the symbiosis alters the plant responsiveness to these signals. OG treatment was previously shown to trigger local and systemic defense responses in tomato plants, involving JA and ET signalling pathways and leading to enhanced resistance to *Botrytis cinerea* (Gamir et al., 2020). Here we show that mycorrhizal tomato plants display primed responses to these signals, and we hypothesise that this may contribute to the enhanced resistance in mycorrhizal plants. We show that the low dose (5 ug/ml) induces significant changes in the proteomic profile of mycorrhizal plants, but not in the non-mycorrhizal ones, that needs a higher dose (50 ug/ml) for a similar activation of defenses. Our work demonstrates for the first time that self-damage perception triggers primed responses in mycorrhizal plants. Previous studies showed that the simultaneous perception of endogenous (DAMPs) and exogenous signals, such as pathogen-associated molecular patterns (PAMPs), primes callose accumulation and PAMP-triggered immunity (PTI) leading to pathogen resistance (Pastor et al., 2022). Interestingly, a possible link between defense priming and self-damage perception has also been suggested in mammals (Krombach et al., 2019). In that study, the perception of protein DAMPs primed immune responses against tumour cells, suggesting a potential correlation between damage recognition and defense priming. In the same line, we propose that the primed responsiveness to self-damage signals in mycorrhizal plants mediates the primed response to pathogens.

The functional description of the differentially accumulated proteins in mycorrhizal plants upon OG elicitation revealed primed synthesis of plant defense related cell wall hydrolases, such as pectinesterases, glucanases and endochitinases, and the proteinase inhibitor Pin2. It is described that some pectinesterases contribute to plant immunity by delivering cell wall-derived signals to the apoplast (Lionetti et al., 2017; Bethke et al., 2014). Inhibitor proteins control pectinesterase activity in plants and, interestingly, oligogalacturonides regulate the activity of the pectinesterase inhibitor, AtPMEI11 (Lionetti et al., 2017). Therefore, it is tempting to speculate that the production of pectinesterases in mycorrhizal plants delivers cell wall fragments, such as oligogalacturonides, in the apoplast, contributing to the primed responses. We further showed that mycorrhiza primes a glucanase biosynthesis in response to OGs. Glucanase activity is a primary defense response against pathogenic microbes (Sela-Buurlage et al., 1993; van Loon, 2006) and, microbial-IR primes glucanase enzyme activity against virulent pathogens (Heil and Bostock, 2002; Siddaiah et al., 2017; Singh and Jha, 2017). It was previously shown that mycorrhizal tomato showing enhanced resistance to the root pathogen *Phytophthora parasitica* displayed higher glucanase activity (Pozo et al., 2002). Furthermore, cucumber plants inoculated with *Trichoderma asperellum* T203 showed that JA and ET signalling pathways regulate systemic glucanase activity in cucumber leaves infected with *Pseudomonas syringae* (Shoresh et al., 2005). Thus, it is plausible that differential glucanase accumulation in mycorrhizal plants is an immune response after the perception of danger signals. Moreover, our results show also a primed accumulation of endochitinase in mycorrhizal plants after OGs treatment. Along with glucanases, chitinases are well known defense related hydrolases which are induced during the plant response to pathogen infection resulting in increased plant resistance (van Loon et al, 2006). Finally, we observed a primed accumulation of Pin2 in mycorrhizal plants in response to OGs. Pin2 is well known wound and JA inducible defense protein that increase plant resistance against herbivorous insects and pathogens (Koiwa et al., 1997). Summarizing, our data show that mycorrhiza primes the biosynthesis of several PR proteins in response to OGs perception, thus preparing the plant to mount a more efficient defense to a broad spectrum of aggressors such as pathogens and insects.

Some of the primed proteins are known to be JA responsive, and the transcriptional analysis also confirmed that AMF colonization primes JA signalling and responsive genes upon OGs recognition. MIR have been shown to be dependent on JA signalling (Fiorilli et al., 2024). Indeed, JA levels are primed in mycorrhizal plants in response to pathogens as *B. cinerea* and to herbivores (Sanchez-Bel et al., 2016, Lidoy et al., 2024), and JA deficient tomato and bean mutants do not show priming nor mycorrhiza induced resistance (Song et al., 2013; Song et al., 2015, Mora-Romero et al., 2014). Previously, we showed that OGs increase JA and ET signalling in tomato plants (Gamir et al., 2020). We suggest that the primed self-damage perception in mycorrhizal plants leads to priming of JA signalling and the induced resistance against pathogens. Remarkably, we found that ET biosynthetic and signalling genes also exhibited primed expression in mycorrhizal plants upon OG elicitation. ET signalling has been suggested as a relevant contributor for microbial-IR and DAMPs-triggered immunity (Shoresh et al., 2005; Liu et al., 2013; Gamir et al., 2020; Lidoy et al., 2024) and has been shown to be required for the primed JA burst leading to MIR in tomato plants (Lidoy et al., 2024). Together, these data suggest a potential role of the JA/ET signalling after damage perception in mycorrhizal plants. Future work is required to clarify the downstream regulation of the JA/ET signalling pathway and the role of the secondary metabolism after self-damage perception in mycorrhiza associated defense priming.

When addressing the changes at the metabolic levels, compound identification of primed signals showed that damage perception in mycorrhizal tomato plants leads to primed flavonoid biosynthesis. Previously we showed that OG application in tomato leaves induces local and systemic biosynthesis of flavonoids (Gamir et al., 2020). Flavonoids are secondary metabolites that provide health benefits and antioxidant properties to the plants, and accordingly, the mycorrhizal status may potentiate the benefits of the OG treatment at this regard. Other beneficial soil microbes as the bacteria *Bacillus subtilis* have been shown to increase the biosynthesis of flavonoids in tomato (Pretali et al., 2016). Our work suggests a potential implication of flavonoids and the plant’s antioxidant system in MIR.

Besides the priming of the above mentioned defensive proteins and compounds, our proteomic analysis revealed the primed accumulation of two receptor kinases in mycorrhizal plants in response to OGs. Extracellular receptor kinases are involved in the perception of endogenous or exogenous signals (Bender and Zipfel, 2023) and damage perception upregulates pattern-recognition receptors in adjacent cells, demonstrating that plants require self-damage and microbial recognition to mount an effective immune response (Zhou et al., 2020). Thus, the enhancement in extracellular receptor kinases can mediate the enhanced responsiveness to OGs observed in mycorrhizal plants. To this regard, we addressed the expression levels of the wak1 gene, and we found that mycorrhizal plants displayed increased wak 1 gene expression. Local upregulation of other wall associated kinases have been recently reported in mycorrhizal cotton roots, regulating root immunity and control of the AM symbiosis (Zang et al, 2024).

Higher levels of receptors is one of the proposed molecular mechanism underlying defense priming, as higher receptor levels will allow faster detection and response to the corresponding signals (Conrath et al., 2015). Similarly, AMF colonization has been shown to trigger the induction of LYS11, an oligosaccharide receptor kinase, in *Lotus japonicus* (Rasmussen et al., 2016) and the production of more oligosaccharide receptors in different plants species (Feng et al., 2019; Girardin et al., 2019). It is therefore tempting to speculate that mycorrhiza colonization upregulates extracellular pattern-recognition receptors, as wak1, enhancing the sensitivity of the plant to changes and consequently priming the plant defensive capacity.

An important question is whether mycorrhizal plants need to perceive damage for a functional MIR. Here, we show that OG recognition is a significant feature for MIR, since MIR against the necrotrophic pathogen *B. cinerea* is abolished in the *wak1* mutant. We show that the tomato mutant *wak1* accumulates less hydrogen peroxide than wild type tomato plants in response to OGs. Further, we demonstrate that OG-IR against *B. cinerea* is abolished in *wak1* plants, confirming that OG perception in the *wak1* mutant is at least partially impaired. In Arabidopsis the WAK family consists of five genes, from WAK1 to WAK5 (He et al., 1999). However, only WAK1 acts as the extracellular receptor of oligogalacturonides (Brutus et al., 2010). Oligogalacturonides treatments induce hydrogen peroxide production and enhance disease resistance against different pathogens in Arabidopsis wild type plants (Galletti et al 2008), and transgenic plants overexpressing WAK1 show increased resistance against several pathogens (Brutus et al., 2010). Not excluding a possible redundant effect of other WAKs in the tomato mutant *wak1*, our data indicate that tomato *wak1* is required for perceiving oligogalacturonides efficiently and for MIR. In tomato, WAK1 acts in a complex with the pattern recognition receptors, Fls2/Fls3, to promote immune responses against the hemibiotrophic bacteria *Pseudomonas syringae* (Zhang et al., 2020). Thus, we hypothesised that the perception of DAMPs, such as oligogalacturonides, promotes not only DAMPs-triggered immunity but also to the primed responses in plants colonised by beneficial microbes.

In conclusion, this study shows that mycorrhizal plants show primed self-damage recognition, and we propose that this feature mediates the primed defense responses to necrotrophic pathogens during MIR. These findings open the “black box” of useful information for a better understanding of the early phases of the immune responses leading to microbial-IR. A similar observation has been made in animal cells, where anti-tumour immunity is dependent on DAMPs recognition, suggesting that early damage recognition could be a general feature for priming responses (Krombach et al., 2019). Interactions between beneficial microbes and plant roots provide plants with an extra layer of defense against pathogens, and the emerging evidences on the role of damage perception in the process deserves further research. Understanding the molecular mechanisms leading to defense priming will contribute to the development of environmentally friendly strategies for crop protection based on the interaction of the plant’s roots and beneficial microbes.

## Abbreviations

AMF: Arbuscular mycorrhizal fungi
AM: Arbuscular mycorrhiza
DAMPs: Damage-Associated Molecular Patterns
ET: Ethylene
Fm: Funneliformis mosseae
IR: Induced resistance
JA: Jasmonic acid
MIR: Mycorrhiza-induced resistance
Nm: Non-mycorrhizal
Ogs: Oligogalacturonides
ROS: Reactive oxygen species
WAKs: Wall-associated kinases

## Supplementary data

Table S1: Table showing the primers used for qPCR analyses.

Table S2: Metabolic pathway annotation of OG-primed metabolites in Funneliformis mosseae mycorrhizal plants

Fig S1: Functional categorization of proteins differentially accumulated in leaves from mycorrhizal tomato.

Fig S2: Functional categorization of proteins from tomato leaves treated with 50 µg/mL of OGs.

## Acknowledgements

We thank Giulia de Lorenzo (Università di Roma “La Sapienza”) for kindly providing the purified oligogalacturonides. We acknowledge the Scientific Instrumentation Service (SCIC) at Universitat Jaume I for their technical assistance, and the proteomic facility PROTEORED© of the University of Alicante (Spain).

## Author Contributions

MJ-P, and JG: conceptualization; ZM, JM-G, EP, and JG: methodology; ZM, and JG: formal analysis; ZM: data curation; JG: writing – original draft; ZM, MJ-P, and JG: writing – review & editing; MJ-P, and JG: supervision; MJ-P, and JG: funding acquisition.

## Data Availability

The datasets generated during the current study are available from the corresponding author on reasonable request.

## References

Adolfsson L, Nziengui H, Abreu IN, Šimura J, Beebo A, Herdean A, Aboalizadeh J, Široká J, Moritz T, Novák O, et al. 2017. Enhanced secondary- and hormone metabolism in leaves of arbuscular mycorrhizal Medicago truncatula. Plant Physiology 175: 392–411.

Bacete L, Mélida H, Miedes E, Molina A. 2018. Plant cell wall-mediated immunity: cell wall changes trigger disease resistance responses. The Plant Journal 93: 614–636.

Basso V, Veneault-Fourrey C. 2020. Role of jasmonates in beneficial microbe–root interactions. In: Champion, A., Laplaze, L. (eds) Jasmonate in Plant Biology. Methods in Molecular Biology, vol 2085. Humana, New York, NY. 43–67. 10.1007/978-1-0716-0142-6_4

Bellincampi D, Dipierro N, Salvi G, Cervone F, De Lorenzo G. 2000. Extracellular H2O2 induced by oligogalacturonides is not involved in the inhibition of the auxin-regulated rolB gene expression in tobacco leaf explants. Plant physiology 122: 1379–1385.

Bender KW, Zipfel C. 2023. Paradigms of receptor kinase signaling in plants. Biochemical Journal 480: 835–854.

Benedetti, M., Mattei, B., Pontiggia, D., Salvi, G., Savatin, D.V., Ferrari, S. 2017. Methods of Isolation and Characterization of Oligogalacturonide Elicitors. In: Shan, L., He, P. (eds) Plant Pattern Recognition Receptors. Methods in Molecular Biology, vol 1578. Humana Press, New York, NY. 10.1007/978-1-4939-6859-6_3

Bethke G, Grundman RE, Sreekanta S, Truman W, Katagiri F, Glazebrook J. 2014. Arabidopsis PECTIN METHYLESTERASEs contribute to immunity against Pseudomonas syringae. Plant Physiology 164: 1093–1107.

Brutus A, Sicilia F, Macone A, Cervone F, De Lorenzo G. 2010. A domain swap approach reveals a role of the plant wall-associated kinase 1 (WAK1) as a receptor of oligogalacturonides. Proceedings of the National Academy of Sciences of the United States of America 107: 9452–9457.

Cameron DD, Neal AL, van Wees SCM, Ton J. 2013. Mycorrhiza-induced resistance: more than the sum of its parts? Trends in Plant Science 18: 539–545.

Chong J, Soufan O, Li C, Caraus I, Li S, Bourque G, Wishart DS, Xia J. 2018. MetaboAnalyst 4.0: towards more transparent and integrative metabolomics analysis. Nucleic Acids Research 46: W486–W494.

Conrath U, Beckers GJM, Langenbach CJG, Jaskiewicz MR. 2015. Priming for Enhanced Defense. Annual Review of Phytopathology 53: 97–119.

Duran-Flores D, Heil M. 2017. Extracellular self-DNA as a damage-associated molecular pattern (DAMP) that triggers self-specific immunity induction in plants. Brain, Behavior, and Immunity.

Edlinger A, Garland G, Hartman K, Banerjee S, Degrune F, García-Palacios P, Hallin S, Valzano-Held A, Herzog C, Jansa J, et al. 2022. Agricultural management and pesticide use reduce the functioning of beneficial plant symbionts. Nature Ecology & Evolution 6: 1145–1154.

Van der Ent S, Van Hulten M, Pozo M, Czechowski T, Udvardi M, Pieterse C, Ton J. 2009. Priming of plant innate immunity by rhizobacteria and beta-aminobutyric acid: differences and similarities in regulation. The New phytologist 183: 419–31.

Feng F, Sun J, Radhakrishnan G V., Lee T, Bozsóki Z, Fort S, Gavrin A, Gysel K, Thygesen MB, Andersen KR, et al. 2019. A combination of chitooligosaccharide and lipochitooligosaccharide recognition promotes arbuscular mycorrhizal associations in Medicago truncatula. Nature Communications 10: 5047.

Ferrari S, Savatin D V, Sicilia F, Gramegna G, Cervone F, Lorenzo G De. 2013. Oligogalacturonides : plant damage-associated molecular patterns and regulators of growth and development. Frontiers in Plant Science 4: 1–9.

Fiorilli V, Martínez-Medina A, Pozo MJ, Lanfranco L. 2024. Plant immunity modulation in arbuscular mycorrhizal symbiosis and its impact on pathogens and pests Annual Review in Phytopathology 62. 10.1146/annurev-phyto-121423-042014

Flors V, Kyndt T, Mauch Mani B, Pozo MJ, Ryu CM, Ton J. 2024. Enabling sustainable crop protection with induced resistance in plants. Frontiers in Science. In press.

Galletti R, Denoux C, Gambetta S, Dewdney J, Ausubel FM, De Lorenzo G, Ferrari S. 2008. The AtrbohD-mediated oxidative burst elicited by oligogalacturonides in Arabidopsis Is dispensable for the activation of defense responses effective against Botrytis cinerea. Plant Physiology 148: 1695–1706.

Gamir J, Minchev Z, Berrio E, García JM, De Lorenzo G, Pozo MJ. 2020. Roots drive oligogalacturonide-induced systemic immunity in tomato. Plant Cell and Environment 44: 275–289.

Gamir J, Pastor V, Kaever A, Cerezo M, Flors V. 2014. Targeting novel chemical and constitutive primed metabolites against Plectosphaerella cucumerina. Plant Journal 78: 227–240.

García JM, Pozo MJ, López-Ráez JA. 2020. Histochemical and molecular quantification of arbuscular mycorrhiza symbiosis. In: Rodríguez-Concepción M, Welsch R, eds. Plant and food carotenoids. Methods in molecular biology. Humana, New York, NY, 293–299.

Giovannetti M, Mosse B. 1980. An evaluation of techniques for measuring vesicular arbuscular mycorrhizal infection in roots. New Phytologist 84: 489–500.

Girardin A, Wang T, Ding Y, Keller J, Buendia L, Gaston M, Ribeyre C, Gasciolli V, Auriac M-C, Vernié T, et al. 2019. LCO receptors involved in arbuscular mycorrhiza are functional for rhizobia perception in legumes. Current Biology 29: 4249–4259.e5.

Gramegna G, Modesti V, Savatin D V., Sicilia F, Cervone F, De Lorenzo G. 2016. GRP-3 and KAPP, encoding interactors of WAK1, negatively affect defense responses induced by oligogalacturonides and local response to wounding. Journal of Experimental Botany 67: 1715–1729.

Gravino M, Savatin DV, Macone A, De Lorenzo G. 2015. Ethylene production in Botrytis cinerea- and oligogalacturonide-induced immunity requires calcium-dependent protein kinases. The Plant Journal 84: 1073–1086.

Gust AA, Pruitt R, Nürnberger T. 2017. Sensing danger: key to activating plant immunity. Trends in Plant Science 22: 779–791.

He Z-H, Cheeseman I, He D, Kohorn BD. 1999. A cluster of five cell wall-associated receptor kinase genes, Wak1-5, are expressed in specific organs of Arabidopsis. Plant Molecular Biology 39: 1189–1196.

He L, Li C, Liu R. 2017. Indirect interactions between arbuscular mycorrhizal fungi and Spodoptera exigua alter photosynthesis and plant endogenous hormones. Mycorrhiza 27: 525–535.

Heil M, Bostock RM. 2002. Induced systemic resistance (ISR) against pathogens in the context of induced plant defences. Annals of Botany 89: 503–512.

Hewitt EJ. 1966. Sand and water culture methods used in the study of plant nutrition. Farnham Royal, England : Commonwealth Agricultural Bureaux.

Hill EM, Robinson LA, Abdul-Sada A, Vanbergen AJ, Hodge A, Hartley SE. 2018. Arbuscular mycorrhizal fungi and plant chemical defence: effects of colonisation on aboveground and belowground metabolomes. Journal of Chemical Ecology 44: 198–208.

van Hulten M, Pelser M, van Loon LC, Pieterse CMJ, Ton J. 2006. Costs and benefits of priming for defense in Arabidopsis. Proceedings of the National Academy of Sciences 103: 5602–5607.

Jewell JB, Sowders JM, He R, Willis MA, Gang DR, Tanaka K. 2019. Extracellular ATP Shapes a Defense-Related Transcriptome Both Independently and along with Other Defense Signaling Pathways. Plant Physiology 179: 1144–1158.

Jones JDG, Staskawicz BJ, Dangl JL. 2024. The plant immune system: From discovery to deployment. Cell 187: 2095–2116.

Jung SC, Martinez-Medina A, Lopez-Raez JA, Pozo MJ. 2012. Mycorrhiza-induced resistance and priming of plant defenses. Journal of Chemical Ecology 38: 651–664.

Kaever A, Landesfeind M, Feussner K, Mosblech A, Heilmann I, Morgenstern B, Feussner I, Meinicke P. 2015. MarVis-Pathway: integrative and exploratory pathway analysis of non-targeted metabolomics data. Metabolomics 11: 764–777.

De Kesel J, Conrath U, Flors V, Luna E, Mageroy MH, Mauch-Mani B, Pastor V, Pozo MJ, Pieterse CMJ, Ton J, et al. 2021. The Induced Resistance Lexicon: Do’s and Don’ts. Trends in Plant Science 26: 685–691.

Kohorn BD. 2016. Cell wall-associated kinases and pectin perception. Journal of Experimental Botany 67: 489–494.

Koiwa H, Bressan RA, Hasegawa PM. 1997. Regulation of protease inhibitors and plant defense. Trends in Plant Science 2: 379–384.

Krombach J, Hennel R, Brix N, Orth M, Schoetz U, Ernst A, Schuster J, Zuchtriegel G, Reichel CA, Bierschenk S, et al. 2019. Priming anti-tumor immunity by radiotherapy: dying tumor cell-derived DAMPs trigger endothelial cell activation and recruitment of myeloid cells. OncoImmunology 8.

Lidoy J, Rivero J, Ramšak Ž, Petek M, Križnik M, Flors V, Lopez-Raez JA, Martinez-Medina A, Gruden K, Pozo MJ. 2024. Ethylene signaling is essential for mycorrhiza-induced resistance against chewing herbivores in tomato. bioRxiv: 2024.06.13.598897.

Lionetti V, Fabri E, De Caroli M, Hansen AR, Willats WGT, Piro G, Bellincampi D. 2017. Three pectin methylesterase inhibitors protect cell wall integrity for Arabidopsis immunity to Botrytis. Plant Physiology 173: 1844–1863.

Liu Z, Wu Y, Yang F, Zhang Y, Chen S, Xie Q, Tian X, Zhou J-M. 2013. BIK1 interacts with PEPRs to mediate ethylene-induced immunity. Proceedings of the National Academy of Sciences 110: 6205–6210.

Livak KJ, Schmittgen TD. 2001. Analysis of relative gene expression data using real-time quantitative PCR and the 2−ΔΔCT method. Methods 25: 402–408.

van Loon LC, Rep M, Pieterse CMJ. 2006. Significance of inducible defense-related proteins in infected plants. Annual Review of Phytopathology 44: 135–162.

De Lorenzo G, Brutus A, Savatin DV, Sicilia F, Cervone F. 2011. Engineering plant resistance by constructing chimeric receptors that recognize damage-associated molecular patterns (DAMPs). FEBS Letters 585: 1521–1528.

De Lorenzo G, Cervone F. 2022. Plant immunity by damage-associated molecular patterns (DAMPs). Essays in Biochemistry 66: 459–469.

De Lorenzo G, Ferrari S, Cervone F, Okun E. 2018. Extracellular DAMPs in plants and mammals: immunity, tissue damage and repair. Trends in Immunology 39: 937–950.

Luna E, Pastor V, Robert J, Flors V, Mauch-Mani B, Ton J. 2011. Callose Deposition: A Multifaceted Plant Defense Response. Molecular Plant-Microbe Interactions® 24: 183–193.

Manresa-Grao M, Pastor V, Sánchez-Bel P, Cruz A, Cerezo M, Jaques JA, Flors V. 2024. Mycorrhiza-induced resistance in citrus against Tetranychus urticae is plant species dependent and inversely correlated to basal immunity. Pest Management Science 80: 3553–3566.

Martinez-Medina A, Flors V, Heil M, Mauch-Mani B, Pieterse CMJ, Pozo MJ, Ton J, van Dam NM, Conrath U. 2016. Recognizing plant defense priming. Trends in Plant Science 21: 818–822.

Martínez-Medina A, Pescador L, Fernández I, Rodríguez-Serrano M, García JM, Romero-Puertas MC and Pozo, MJ. 2019. Nitric oxide and phytoglobin PHYTOGB1 are regulatory elements in the Solanum lycopersicum–Rhizophagus irregularis mycorrhizal symbiosis. New Phytologist 223: 1560–1574.

Molina A, Jordá L, Torres MÁ, Martín-Dacal M, Berlanga DJ, Fernández-Calvo P, Gómez-Rubio E, Martín-Santamaría S. 2024. Plant cell wall-mediated disease resistance: Current understanding and future perspectives. Molecular Plant 17: 699–724.

Mora-Romero GA, Gonzalez-Ortiz MA, Quiroz-Figueroa F, Calderon-Vazquez CL, Medina-Godoy S, Maldonado-Mendoza I, Arroyo-Becerra A, Perez-Torres A, Alatorre-Cobos F, Sanchez F, et al. 2015. PvLOX2 silencing in common bean roots impairs arbuscular mycorrhiza-induced resistance without affecting symbiosis establishment. Functional Plant Biology 42: 18.

Meco V, Egea I, Ortíz-Atienza A, Drevensek S, Esch E, Yuste-Lisbona FJ, Barneche F, Vriezen W, Bolarin MC, Lozano R, et al. 2020. The salt sensitivity induced by disruption of cell wall-associated kinase 1 (SlWAK1) tomato gene is linked to altered osmotic and metabolic homeostasis. International Journal of Molecular Sciences 21: 6308.

Nair A, Kolet SP, Thulasiram H V., Bhargava S. 2015. Role of methyl jasmonate in the expression of mycorrhizal induced resistance against Fusarium oxysporum in tomato plants. Physiological and Molecular Plant Pathology 92: 139–145.

Pastor V, Cervero R, Gamir J. 2022. The simultaneous perception of self- and non-self-danger signals potentiates plant innate immunity responses. Planta 256: 10.

Pieterse CMJ, Leon-Reyes A, Van der Ent S, Van Wees SCM. 2009. Networking by small-molecule hormones in plant immunity. Nature chemical biology 5: 308–316.

Pieterse CMJ, Zamioudis C, Berendsen RL, Weller DM, Van Wees SCM, Bakker PAHM. 2014. Induced systemic resistance by beneficial microbes. Annual Review of Phytopathology 52: 347–375.

Pontiggia D, Benedetti M, Costantini S, De Lorenzo G, Cervone F. 2020. Dampening the DAMPs: How Plants Maintain the Homeostasis of Cell Wall Molecular Patterns and Avoid Hyper-Immunity. Frontiers in Plant Science 11.

Pozo MJ, Azcón-Aguilar C. 2007. Unraveling mycorrhiza-induced resistance. Current Opinion in Plant Biology 10: 393–398.

Pozo MJ, Cordier C, Dumas-Gaudot E, Gianinazzi S, Barea JM, Azcón-Aguilar C. 2002. Localized versus systemic effect of arbuscular mycorrhizal fungi on defence responses to Phytophthora infection in tomato plants. Journal of Experimental Botany 53: 525–534.

Pozo MJ, López-Ráez JA, Azcón-Aguilar C, García-Garrido JM. 2015. Phytohormones as integrators of environmental signals in the regulation of mycorrhizal symbioses. New Phytologist 205: 1431–1436.

Pozo MJ, Zabalgogeazcoa I, Vazquez de Aldana BR, Martinez-Medina A. 2021. Untapping the potential of plant mycobiomes for applications in agriculture. Current Opinion in Plant Biology 60: 102034.

Pretali L, Bernardo L, Butterfield TS, Trevisan M, Lucini L. 2016. Botanical and biological pesticides elicit a similar Induced Systemic Response in tomato (Solanum lycopersicum) secondary metabolism. Phytochemistry 130: 56–63.

Rasmussen SR, Füchtbauer W, Novero M, Volpe V, Malkov N, Genre A, Bonfante P, Stougaard J, Radutoiu S. 2016. Intraradical colonization by arbuscular mycorrhizal fungi triggers induction of a lipochitooligosaccharide receptor. Scientific Reports 6: 29733.

Rassizadeh L, Cervero R, Flors V, Gamir J. 2021. Extracellular DNA as an elicitor of broad-spectrum resistance in Arabidopsis thaliana. Plant Science 312: 111036.

Rivero J, Álvarez D, Flors V, Azcón-Aguilar C, Pozo MJ. 2018. Root metabolic plasticity underlies functional diversity in mycorrhiza-enhanced stress tolerance in tomato. New Phytologist 220: 1322–1336.

Rivero J, Lidoy J, Llopis-Giménez Á, Herrero S, Flors V, Pozo MJ. 2021. Mycorrhizal symbiosis primes the accumulation of antiherbivore compounds and enhances herbivore mortality in tomato. Journal of Experimental Botany 72: 5038–5050.

Sanchez-Bel P, Troncho P, Gamir J, Pozo MJ, Camañes G, Cerezo M, Flors V. 2016. The nitrogen availability interferes with mycorrhiza-induced resistance against Botrytis cinerea in tomato. Frontiers in Microbiology 7: 1–16.

Sanmartín N, Pastor V, Pastor-Fernández J, Flors V, Pozo MJ, Sánchez-Bel P. 2020a. Role and mechanisms of callose priming in mycorrhiza-induced resistance. Journal of Experimental Botany 71: 2769–2781.

Sanmartín N, Sánchez-Bel P, Pastor V, Pastor-Fernández J, Mateu D, Pozo MJ, Cerezo M, Flors V. 2020b. Root-to-shoot signalling in mycorrhizal tomato plants upon Botrytis cinerea infection. Plant Science 298: 110595.

Schoenherr AP, Rizzo E, Jackson N, Manosalva P, Gomez SK. 2019. Mycorrhiza-induced resistance in potato involves priming of defense responses against cabbage looper (Noctuidae: Lepidoptera). Environmental Entomology 48: 370–381.

Scortica A, Giovannoni M, Scafati V, Angelucci F, Cervone F, De Lorenzo G, Benedetti M, Mattei B. 2022. Berberine Bridge Enzyme-like Oligosaccharide Oxidases Act as Enzymatic Transducers Between Microbial Glycoside Hydrolases and Plant Peroxidases. Molecular Plant-Microbe Interactions® 35: 881–886.

Sela-Buurlage MB, Ponstein AS, Bres-Vloemans SA, Melchers LS, van den Elzen PJM, Cornelissen BJC. 1993. Only specific tobacco (Nicotiana tabacum) chitinases and [beta]-1,3-glucanases exhibit antifungal activity. Plant Physiology 101: 857–863.

Shoresh M, Yedidia I, Chet I. 2005. Involvement of jasmonic acid/ethylene signaling pathway in the systemic resistance induced in cucumber by Trichoderma asperellum T203. Phytopathology® 95: 76–84.

Siddaiah CN, Satyanarayana NR, Mudili V, Kumar Gupta V, Gurunathan S, Rangappa S, Huntrike SS, Srivastava RK. 2017. Elicitation of resistance and associated defense responses in Trichoderma hamatum induced protection against pearl millet downy mildew pathogen. Scientific Reports 7: 43991.

Singh RP, Jha PN. 2017. The PGPR Stenotrophomonas maltophilia SBP-9 augments resistance against biotic and abiotic stress in wheat plants. Frontiers in Microbiology 8.

Smith SE, Smith FA. 2011. Roles of arbuscular mycorrhizas in plant nutrition and growth: New paradigms from cellular to ecosystem scales. Annual Review of Plant Biology 62: 227–250.

Smith CA, Want EJ, O’Maille G, Abagyan R, Siuzdak G. 2006. XCMS: Processing mass spectrometry data for metabolite profiling using nonlinear peak alignment, matching, and identification. Analytical Chemistry 78: 779–787.

Song Y, Chen D, Lu K, Sun Z, Zeng R. 2015. Enhanced tomato disease resistance primed by arbuscular mycorrhizal fungus. Frontiers in Plant Science 6.

Song YY, Ye M, Li CY, Wang RL, Wei XC, Luo SM, Zeng R Sen. 2013. Priming of Anti-Herbivore Defense in Tomato by Arbuscular Mycorrhizal Fungus and Involvement of the Jasmonate Pathway. Journal of Chemical Ecology 39: 1036–1044.

Souza C de A, Li S, Lin AZ, Boutrot F, Grossmann G, Zipfel C, Somerville SC. 2017. Cellulose-derived oligomers act as damage-associated molecular patterns and trigger defense-like responses. Plant Physiology 173: 2383–2398.

Stephens C, Hammond-Kosack KE, Kanyuka K. 2022. WAKsing plant immunity, waning diseases. Journal of Experimental Botany 73: 22–37.

Tanaka K, Heil M. 2021. Damage-Associated Molecular Patterns (DAMPs) in Plant Innate Immunity: Applying the Danger Model and Evolutionary Perspectives. Annual Review of Phytopathology 59: 53–75.

Tripathi D, Zhang T, Koo AJ, Stacey G, Tanaka K. 2018. Extracellular ATP acts on jasmonate signaling to reinforce plant defense. Plant Physiology 176: 511–523.

Velivelli SL, Lojan P, Cranenbrouck S, de Boulois HD, Suarez JP, Declerck S, Franco J, Prestwich BD. 2015. The induction of Ethylene response factor 3 (ERF3) in potato as a result of co-inoculation with Pseudomonas sp. R41805 and Rhizophagus irregularis MUCL 41833 – a possible role in plant defense. Plant Signaling & Behavior 10: e988076.

Wagner TA, Kohorn BD. 2001. Wall-Associated kinases are expressed throughout plant development and are required for cell expansion. The Plant Cell 13: 303–318.

Wu C-T, Bradford KJ. 2003. Class I chitinase and β-1,3-glucanase are differentially regulated by wounding, methyl jasmonate, ethylene, and gibberellin in tomato seeds and leaves. Plant Physiology 133: 263–273.

Zhang M, Kong X. 2022. How plants discern friends from foes. Trends in Plant Science 27: 107–109.

Zhang N, Pombo MA, Rosli HG, Martin GB. 2020. Tomato wall-associated kinase SlWak1 depends on Fls2/Fls3 to promote apoplastic immune responses to Pseudomonas syringae. Plant Physiology 183: 1869–1882.

Zhang X, Jia S, He Y, Wen J, Li D, Yang W, Yue Y, Li H, Cheng K, Zhang X. 2024. Wall-associated kinase GhWAK13 mediates arbuscular mycorrhizal symbiosis and Verticillium wilt resistance in cotton. New Phytologist 242: 2180–2194.

Zhou F, Emonet A, Dénervaud Tendon V, Marhavy P, Wu D, Lahaye T, Geldner N. 2020. Co-incidence of damage and microbial patterns controls localized immune responses in roots. Cell 180: 440–453.e18.

Zipfel C, Oldroyd GED. 2017. Plant signalling in symbiosis and immunity. Nature 543: 328–336.

